# CaveCrawler: An interactive analysis suite for cavefish bioinformatics

**DOI:** 10.1101/2021.12.01.470856

**Authors:** Annabel Perry, Suzanne E. McGaugh, Alex C. Keene, Heath Blackmon

## Abstract

The growing use of genomics data in diverse animal models provides the basis for identifying genomic and transcriptional differences across species and contexts. Databases containing genomic and functional data have played critical roles in the development of numerous genetic models but are lacking for most emerging models of evolution. There is a rapidly expanding use of genomic, transcriptional, and functional genetic approaches to study diverse traits of the Mexican tetra, *Astyanax mexicanus*. This species exists as two morphs, eyed surface populations and at least 30 blind cave populations, providing a system to study convergent evolution. We have generated a web-based analysis suite that integrates datasets from different studies to identify how gene transcription and genetic markers of selection differ between populations and across experimental contexts. Results can be processed with other analysis platforms including Gene Ontology (GO) to enable biological inference from cross-study patterns and identify future avenues of research. Furthermore, the framework that we have built *A. mexicanus* can readily applied to other emerging model systems.

## Introduction

The reduced cost and increased efficiency of sequencing has led to enormous growth in the application of sequencing approaches to study diverse biological processes. In previous decades, these approaches were predominantly performed on a small number of genetically amendable model organisms including *Caenorhabditis elegans, Drosophila melanogaster*, zebrafish, and mouse. Model-organism-specific databases have been generated for each of these model systems, providing critical resources that decrease access barriers to genomic and phenotypic data (1–3). Recently, there has been increased application of genomic and molecular approaches to non-standard model systems, as these model systems may enable comparative evolutionary studies not possible in traditional systems (4). However, a lack of databases and analytic tools for many of these emerging model organisms impedes analysis of genomic data collected across different studies.

The Mexican tetra, *Astyanax mexicanus* is an emerging model system to study the convergent evolution of diverse biological traits. These fish are comprised of a single population of river dwelling surface fish and at least 30 cavefish populations of the same species (5). *Astyanax mexicanus* cavefish populations have independently evolved numerous morphological, behavioral, and physiological differences from their surface conspecifics (6,7). These fish can be efficiently reared in laboratories, allowing for the application of transgenic and gene-editing approaches (8). There is a rapidly growing focus on genomic data in these systems that compare cave and surface populations. Current genomic data includes fully assembled genomes for surface and cave populations, population genetic resequencing, and transcriptomic data across different contexts (9,10). The development of a database that compiles the growing number of genomics data across different contexts would provide a valuable resource for accessing and analyzing this information.

The Shiny package in R offers a method to produce powerful community web resources that can go far beyond traditional repositories of data (11). Shiny databases enable researchers to incorporate the statistical analysis and data visualization capabilities of the R programming language into a reactive database that also functions as a community data repository. The combination of these tools allows users to sift through vast amounts of data, enabling novel discoveries (12). The generation of a Shiny database for comparative models of evolution could combine data across populations and studies. The flexibility of these systems and intrinsic analysis capabilities allows for direct comparisons of genetic data from disparate sources. Here, we generated a Shiny database, CaveCrawler, which combines population genetics and transcriptomic data from multiple Mexican tetra populations and leverages Gene Ontology (GO) term information to enable unique biological inferences from cross-study patterns. We demonstrate that the analysis features of this program can identify genes that are implicated in evolutionary processes across populations of *A. mexicanus*, using different methodologies, and in different studies.

## Methods

The CaveCrawler database acts as a repository for transcription, Gene Ontology (GO), population genetics, and annotated genome data acquired from different studies in *A. mexicanus*, including those using reference genomes for surface and Pachón cavefish (9,13). With a highly accessible web interface, CaveCrawler enables researchers to search for data on genes-of-interest, find genes whose transcriptional levels match defined criteria, find genes which fit desired population genetics parameters, and also identify genes associated with cellular components, molecular functions, and biological processes.

### CaveCrawler modules

The CaveCrawler framework utilizes a bifurcated design with an underlying data repository and a collection of user interface modules (Figure 1). The databases currently offers five user modules: Home, Gene Search, Transcription, Population Genetics, and GO Term Info. Each of these modules is designed to draw on different elements of the underlying data repository. This bifurcated design facilitates simple updates to the repository which then are immediately populated into changes in the functionality and results produced by the modules that draw on the updated repositories. Similarly, new modules can be added at any time to take advantage of new types of analyses users desire or new data types included in the repository. The home module houses general information about *A. mexicanus* and about CaveCrawler’s functionality, as well brief instructions for contributing data.

**Figure 1.**
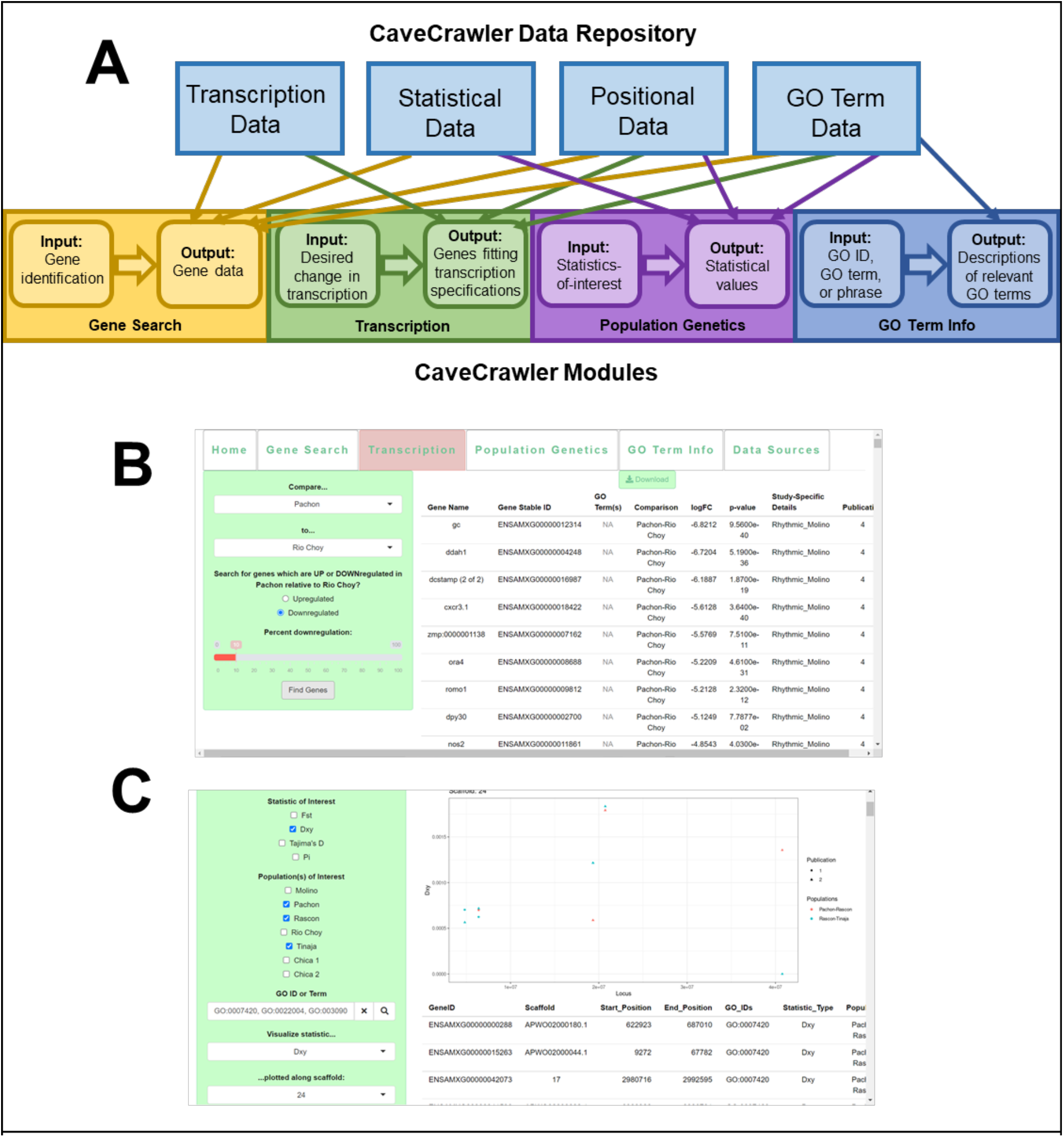
Design and web interface for CaveCrawler. A) The repository and module framework for the CaveCrawler model organism genomics database. Lines show the connections between different types of data stored in the repository and the user modules that draw on each data type. B) Example of the Transcription module with the results of searching for top 10% of genes that are downregulated in Pachón relative to Rio Choy surface fish C) Example of the Population Genetics module with the results of searching for the Pachón-Rascon surface fish and Rascon-Tinaja d_XY_ values of genes associated with brain development GO IDs and visualizing these values on Scaffold 24

The Gene Search module enables the user to search for data associated with genes-of-interest and also to identify genes associated with GO terms-of-interest. In this module, the user inputs a single gene stable ID, a single GO term, or a comma-separated list of genes. The module outputs a downloadable table describing all genes associated with the inputs and the positional, transcription, and population genetics data associated with each of the genes. The output also indicates whether a statistic or piece of transcriptional data is not present for each gene-of-interest. Therefore, this module concatenates data from disparate sources into a single analysis output, enabling the user to efficiently search for existing data and identify experiments which have yet to be conducted on their genes-of-interest.

The Transcription module enables the user to identify genes which differ in transcription level between groups. Here, the user first inputs the groups they would like to compare. The user may either compare an experimental group to a control group or compare one morph to another morph. The user then specifies whether they would like to see genes which are up or downregulated in the first group compared to the second and the percent change in transcription level between groups. The module then produces a downloadable output table of genes fitting the specified transcription patterns.

The Population Genetics module enables the user to access population genomics statistics, such as π, Tajima’s D, d_XY_, and F_ST_. This module has two options for accessing population genomics data. In the first option, the user provides GO terms and the module outputs and visualizes the statistical values of all genes associated with those GO terms. The second approach enables the user to search for transcriptional or genomic values associated with defined across different analyses.

In the GO term search function of the Population Genetics module, the user inputs GO information, statistics-of-interest, and populations-of-interest. For the GO information, the user can input either a single GO ID, a comma-separated list of GO IDs, or a phrase associated with the target GO term. The module outputs a downloadable table describing all values of the population-specific statistics-of-interest for the genes associated with the indicated GO term(s). If any of the statistics-of-interest require pairwise comparisons between populations, the module will output pairwise statistics for each possible pairing of input populations. On this submodule, the user may also input a statistic and a scaffold and CaveCrawler will plot the statistical values of each GO-term-associated gene which falls on that scaffold. The GO term function of the Population Genetics module thus enables the user to access and visualize population genomics statistics for a GO term of interest.

The outlier function of the Population Genetics module consists of two approaches for pulling outlier genes from combined datasets. One approach enables the user to identify a specified number of genes which have the most extreme values for an indicated statistic, while the other approach enables the user to identify all genes whose statistic value falls above or below a specified threshold value. In the gene number approach, the user must specify the number of genes and must specify whether they would like to see the top or bottom quantile. CaveCrawler then outputs a table describing the specified number of genes with the most extreme values for the statistic-of-interest. In the statistical threshold approach, the user specifies a threshold statistical value and specifies whether they would like to see genes above or below this value. CaveCrawler outputs both a table and a distribution plot describing the genes which fall above or below this threshold.

Both outlier approaches require the user input a statistic-of-interest and population(s)-of-interest. If the statistic-of-interest is a one-population statistic, such as π or Tajima’s D, both approaches will report outlier statistical values for all input populations. If the input statistic is a pairwise statistic, such as F_ST_ or d_XY_, both approaches will report outlier statistical values for all possible pairs of populations-of-interest. If a statistic value has yet to be collected for a population or population pair, CaveCrawler will output a warning about that statistic. Thus, the outlier function of the Population Genetics module enables users to not only identify outliers for a statistic-of-interest but also to identify populations for which a statistic-of-interest has yet to be collected.

The GO Term Info module enables users to access descriptions of GO IDs. This function helps users identify GO IDs they should search for in the Population Genetics module and helps them make sense of transcription and outlier queries. On this module, the user may input a single GO ID, comma-separated list of GO IDs, or a phrase-of-interest, such as “sleep”. CaveCrawler searches data from the official Gene Ontology databank, outputting descriptions of all input GO IDs or GO IDs relevant to the input phrase. In addition, CaveCrawler reports all GO IDs which occur hierarchically beneath these IDs. The GO Term Info module thus enables researchers to investigate the broader biological impact of transcription and diversity data relevant to their genes-of-interest.

### The data repository

CaveCrawler pools data from multiple publications and authors can request that their own data be integrated into CaveCrawler’s repository. As of publication, CaveCrawler’s data bank includes transcriptional datasets (14,15), population genetics datasets (10,15), GO data from UniProt and the Gene Ontology Consortium (16–18), and genome architecture data from Ensembl Genome Browser, release 104 (19).

CaveCrawler’s Transcription and Gene Search modules currently draw upon datasets that describe genes whose transcription levels changed significantly in response to sleep deprivation in *A. mexicanus* (20). This dataset describes the log fold-change (logFC) and p-values for each of these genes in each *A. mexicanus* morph where the change in transcription was significant compared to controls of the same morph (20). As described in the Transcription module section of this paper, CaveCrawler can also access transcription data for genes whose transcription is significantly different between morphs (14). The Transcription module has enough flexibility that new transcriptional data can be integrated. Thus, CaveCrawler could be used to analyze transcriptional changes in response to any experimental condition and between any two morphs of *A. mexicanus*.

CaveCrawler’s Population Genetics and Gene Search modules currently integrate data from two studies describing signatures of selection in *A. mexicanus* (10,15). One of these studies calculated π and Tajima’s D values for the Pachon, Tinaja, Molino, Río Choy, and Rascon populations, as well as F_ST_ and d_XY_ values for each population pair (10). The other study describes d_XY_ values of all genes in two populations of the Chica morph, Pachon and Rascon, and Tinaja and Rascon (15). As with the Transcription module, the Population and Gene Search modules have enough flexibility that new data can be integrated.

The Gene Search, Transcription, and Population Genetics modules currently draw upon positional data obtained from Ensembl (19). The genome assembly used in the current version is *A. mexicanus* 2.0, the most up-to-date genome assembly for this species (9). All of CaveCrawler’s modules utilize GO term information from UniProt and from the Gene Ontology Consortium (16–18).

Though CaveCrawler already integrates data from numerous disparate sources, enabling powerful cross-study comparisons of genetic data, CaveCrawler’s data repository is not static. The CaveCrawler website includes instructions for data submission and the power and insights possible with this resource will grow as the repository of data on which draws grows. CaveCrawler’s data repository will be updated annually in July.

## Results

The CaveCrawler analysis suite consists of multiple tools for comparing datasets that allow for identification of genetic differences between populations of *A. mexicanus*. These tools have a wide range of applications, including rapid candidate gene identification and inference of population-level variation. Here, we present an example of how CaveCrawler can be used to answer biological questions.

### Rapid Identification of Candidate Genes for Empirical Studies

Since CaveCrawler enables simultaneous cross-analysis of multiple studies, researchers can use CaveCrawler to find genes which are outliers for both transcription and population genetics statistics in a matter of minutes. These genes can then be analyzed in downstream studies, such as GO term analyses, to make biological inferences. Here, we identified genes which are transcriptionally dysregulated between cave and Río Choy morphs, then performed a GO term analysis to determine the biological function and cellular components with which these genes are associated. These genes could be used as candidates for future empirical studies, such as knockdown or knockout studies.

To examine genes that are both transcriptionally upregulated and harbor markers of selection, we first used CaveCrawler’s Population Genetics module to identify the F_ST_ values of all genes whose F_ST_ values were published in a recent population inference paper in the Mexican tetra (see 10). Then, we used the ‘Gene Search’ module to identify the transcription data for each of the 1140 genes identified by the Population Genetics module (see Supplemental Table 1). By using the Gene Search module, we found that previous studies had measured the between-morph logFC values for 267 of the 1140 genes for which we had F_ST_ values. Pairwise F_ST_ measures how dissimilar a DNA sequence is between two groups relative to diversity within the groups, and logFC is the log fold change in mRNA transcription between two groups (14,21). Of the genes for which both F_ST_ and logFC had been calculated by previous studies, there were 72 for which F_ST_ outlier status had been determined by a previous study (10). Gene names, logFC, transcription p-values, and F_ST_ values for all 72 genes are available in the supplemental materials.

For each cave-Río Choy surface comparison, we then identified the genes which were both significantly differentially expressed for circadian regulation (logFC p-value < 0.05) between Río Choy and the corresponding cave population and were identified by a previous study to be F_ST_ outliers for the same population pairing (10). These genes, which were both transcriptional and F_ST_ outliers, will henceforth be referred to as double outliers. We found one gene which was a double outlier in all three cave-Río Choy pairings (Table 1; Figure 2), one which was a double outlier for both Pachón-Río Choy and for Tinaja-Río Choy (Table 1; Figure 2A and 2C), one which was a double outlier for Molino-Río Choy only (Table 1; Figure 2B), one which was a double outlier for Tinaja-Río Choy only (Table 1; Figure 2C).

**Figure 2.**
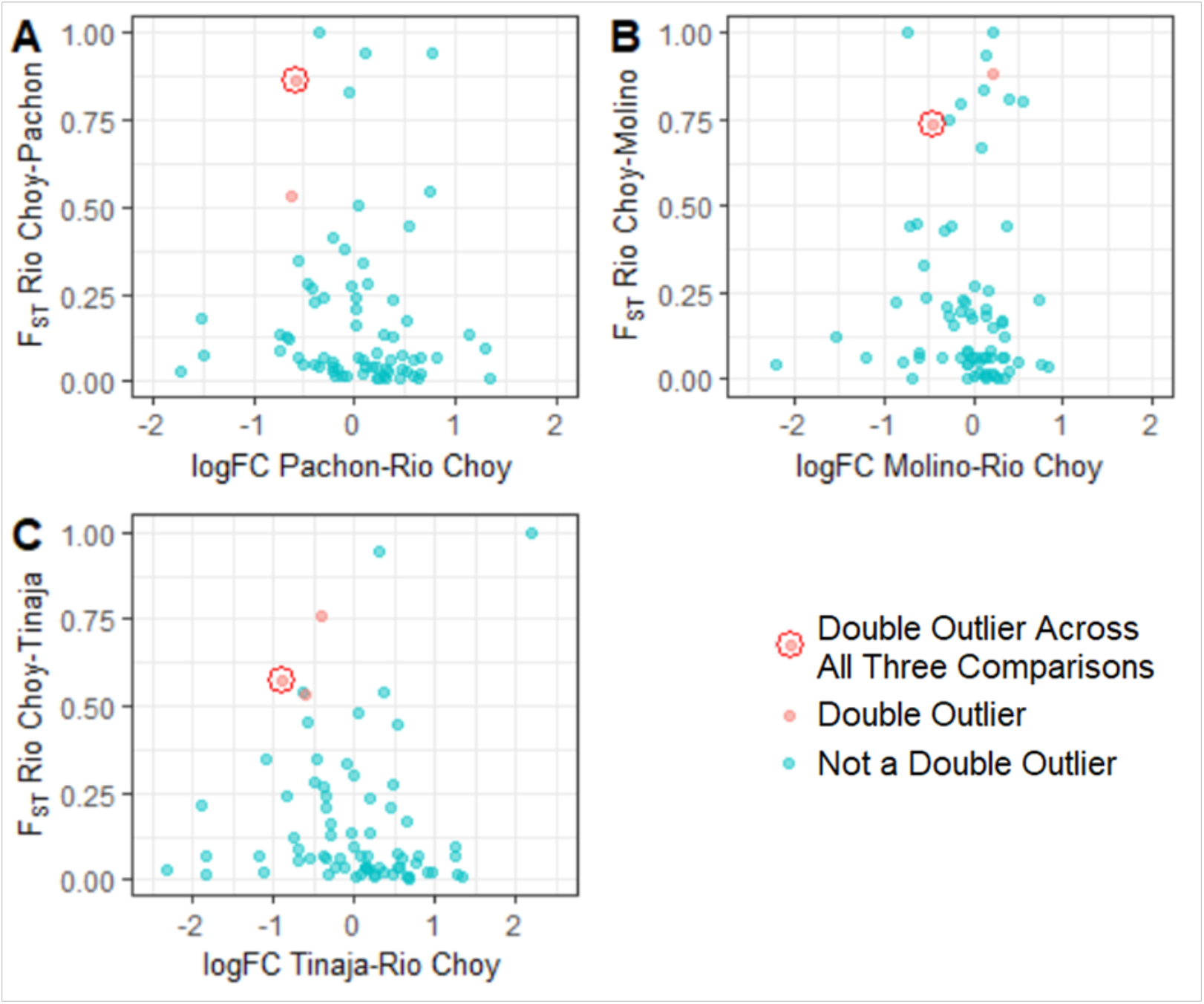
Overlap between F_ST_ values and logFC values across multiple studies. Plots of cave-specific F_ST_ vs. logFC values for all 72 genes which CaveCrawler found to have F_ST_ values, logFC values, and F_ST_ outlier designations. Double-outliers for the cave-Río Choy comparison indicated by the axes are colored in red, while the gene (*arpin*) which was a doubleoutlier across in all 3 cave-Río Choy comparisons is encircled in red. A) Pachón vs. Rio Choy B) Molino vs. Rio Choy C) Tinaja vs. Rio Choy.

**Table 1:**
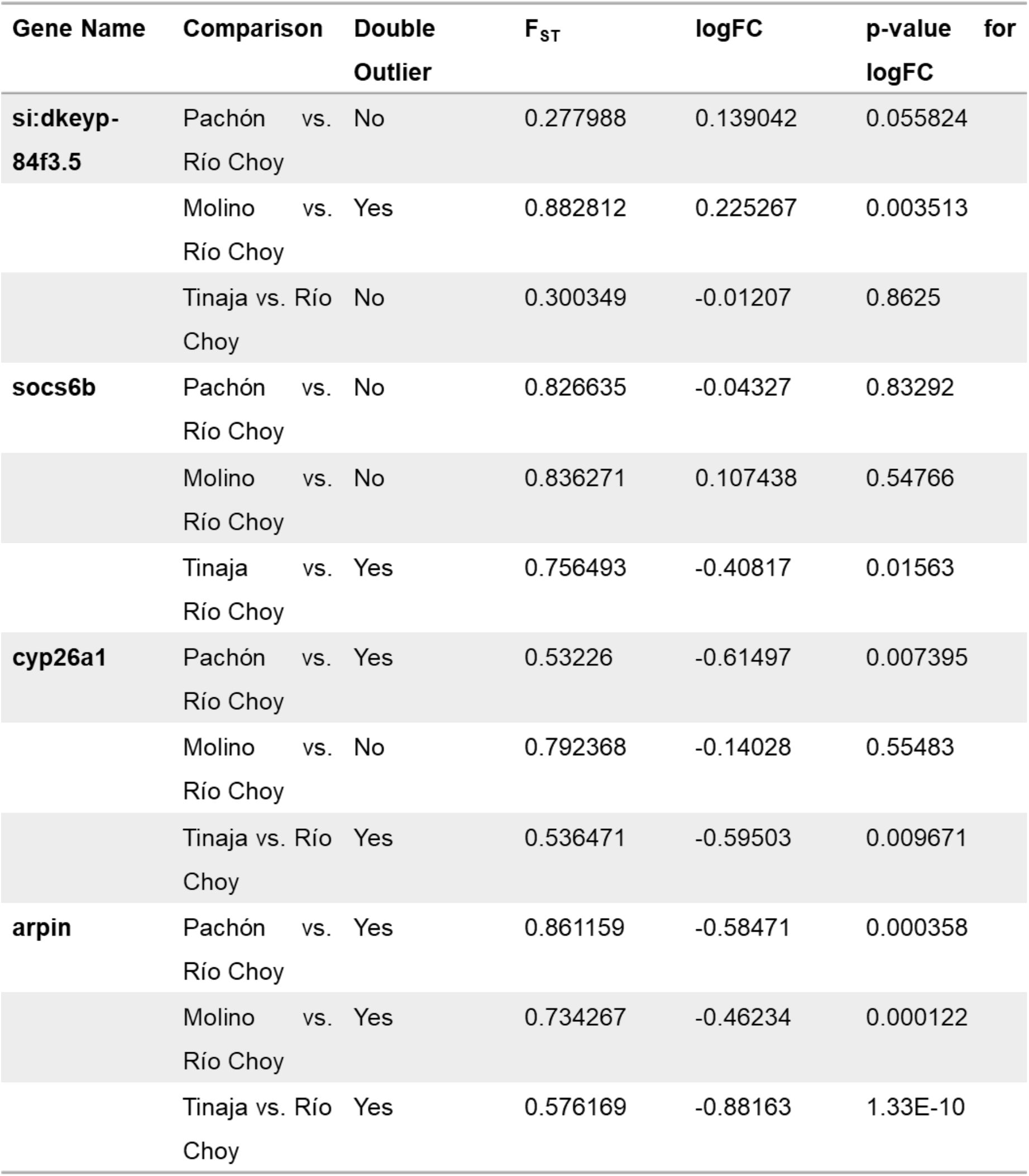
Genes identified as outliers for F_ST_ and transcriptional regulation over the circadian cycle between surface fish and three different cavefish populations. F_ST_ and logFC values for all genes which were found to be outliers for both F_ST_ and logFC in at least one cave-Río Choy comparison

We performed a GO term analysis on *arpin to* identify any biological process, molecular function, or cellular component associated with this double outlier. We found *arpin* to be associated with the biological process GO ID GO:0051126 and the cellular component GO IDs GO:0016021 and GO:0030027, which correspond to “negative regulation of actin nucleation”, “integral component of membrane”, and “lamellipodium”, respectively. To calculate the likelihood of sampling an *A. mexicanus* gene associated with GO:0051126 by chance, we performed a Monte Carlo simulation for 1000000 iterations and calculate an empirical p-value of 2.8e-05. We performed another Monte Carlo to find the likelihood of sampling GO:0016021 and GO:0030027 by chance, obtaining an empirical p-value of 4.4e-05. Thus, we used CaveCrawler to rapidly discover that genes that harbor markers of selection and are transcriptionally in cave populations across the circadian cycle.

As shown by this example, the CaveCrawler analysis suite can be used for a variety of investigations in the Mexican tetra. CaveCrawler can in minutes combine statistics from multiple studies and leverage GO terms to make novel inferences about evolutionary forces acting within a population.

## Discussion

Here, we describe a modular analysis suite for *A. mexicanus*. We have included a set of the genomics and transcriptional data that has been previously published. In addition to these studies, transcriptional analysis across developmental timepoints, as well as single cell analysis of hypothalamus has been collected. These data sets, and others collected in the future can be added to this analysis suite. These data, in combination with assembled genomes for surface fish and Pachón cavefish provide a platform for gene discovery in this system. In addition, the modularity of this system allows it to be readily adapted for new data types or genomic analyses. We then demonstrated that this analysis suite can be used to combine data from disparate sources to discover novel patterns in the Mexican tetra genome.

As proof of principle, we performed an analysis for genes that contained markers of selection and transcriptional dysregulation across the circadian cycle. This analysis identified four genes that were significantly different. These genes represent strong candidate for functional regulators of evolved differences in circadian behavior that have been widely studied in *A. mexicanus* and other species of cavefish (14,22-25). The gene *arpin*, a negative regulator of *actin* is of particular interest because it is identified as harboring markers of selection and transcriptional dysregulation across all three cavefish populations. Actin dynamics have been implicated as targets of circadian regulation for a number of processes including wound healing, immune function and neural plasticity (26-28). Therefore, it is possible that multiple populations of cavefish have converged on changes in actin regulation that account for loss of behavioral and transcriptional rhythms (14,24).

Shiny has been widely applied to develop a range of public databases that offer interactive data visualization and access (12,29,30). However, to our knowledge, this is the first use of Shiny to create a public genomic database and analysis tool for any model organisms. Traditionally these resources which are key to supporting model organism communities have come with considerable cost in the form of computer programmers and hosting services (31,32). Perhaps one of the most valuable contributions that CaveCrawler can make is as a flexible framework that can be adopted by any model organism community. We have made the underlying code for this project publicly available under the GPL license. All source code and example datasets are available in the GitHub repository: https://github.com/AnnabelPerry/AstyanaxShinyApp.

In *A. mexicanus*, like many other models of evolution, studies identifying quantitative trait loci (QTL) have provided a basis for a growing genetic toolkit in *A. mexicanus* can be used for functional genomics experiments guided CaveCrawler (7,33). For example, transgenesis, CRISPR-based transgenesis, and morpholinos have all been applied for functional validation of gene function (34–37). In addition, CRISPR-based screening approaches have been developed in zebrafish that allow for high throughput functional assessment of developmental and behavioral traits. This analysis suite will provide methodology for identifying genes for functional analysis.

## Acknowledgements

This work was supported by an NIH NIGMS R35GM138098 to HB, NIH R01 1R01GM127872 to ACK, and SEM, NIH R21 NS122166 to ACK, and the Texas A&M University College of Science Undergraduate Research Opportunities Program to ARP

